# MeCP2 Interacts with the Super Elongation Complex to Regulate Transcription

**DOI:** 10.1101/2024.06.30.601446

**Authors:** Jun Young Sonn, Wonho Kim, Marta Iwanaszko, Yuki Aoi, Yan Li, Luke Parkitny, Janice L. Brissette, Lorin Weiner, Ismael Al-Ramahi, Juan Botas, Ali Shilatifard, Huda Y. Zoghbi

## Abstract

Loss-of-function mutations in methyl-CpG binding protein 2 (*MECP2*) cause Rett syndrome, a postnatal neurodevelopmental disorder that occurs in ∼1/10,000 live female births. MeCP2 binds to methylated cytosines across genomic DNA and recruits various partners to regulate gene expression. MeCP2 has been shown to repress transcription *in vitro* and interacts with co-repressors such as the Sin3A and NCoR complexes. Based on these observations, MeCP2 has been largely considered as a repressor of transcription. However, a mouse model of RTT displays many down-regulated genes, and those same genes are up-regulated in a *MECP2* duplication mouse model. Furthermore, TCF20, which has been associated with transcriptional activation, have recently been identified as a protein interactor of MeCP2. These data broaden the potential functions of MeCP2 as a regulator of gene expression. Yet, the molecular mechanisms underlying MeCP2-dependent gene regulation remain largely unknown. Here, using a human *MECP2* gain-of-function *Drosophila* model, we screened for genetic modifiers of *MECP2*-induced phenotypes. Our approach identified several subunits of the *Drosophila* super elongation complex, a P-TEFb containing RNA polymerase II (RNA pol II) elongation factor required for the release of promoter-proximally paused RNA pol II, as genetic interactors of *MECP2*. We discovered that MeCP2 physically interacts with the SEC in human cells and in the mouse brain. Furthermore, we found that MeCP2 directly binds AFF4, the scaffold of the SEC, via the transcriptional repression domain. Finally, loss of MeCP2 in the mouse cortex caused reduced binding of AFF4 specifically on a subset of genes involved in the regulation of synaptic function, which also displayed the strongest decrease in RNA pol II binding in the genebody. Taken together, our study reveals a previously unrecognized mechanism through which MeCP2 regulates transcription, providing a new dimension to its regulatory role in gene expression.

## INTRODUCTION

Rett syndrome (RTT) is a devastating neurodevelopmental disorder resulting from *de novo* loss-of-function mutations in the X-linked gene, *MECP2*^1^. Typically, girls with classical RTT develop normally for the first 12-18 months of life, but then lose their acquired language and motor skills and gradually develop stereotypies, motor difficulties, autism, seizures, and autonomic dysfunction^2,3^.

The MeCP2 protein is present in a variety of tissues in mammals, with the highest levels in the neurons of the central nervous system^4,5^. MeCP2 levels are initially low during embryonic development but increases dramatically after birth and is maintained at a high level throughout adulthood^6,7^. The period of MeCP2 increase coincides with the critical stage of postnatal brain development during which neuronal axons grow, synapses form, and experiences refine synaptic connections^8^. Therefore, MeCP2 is believed to be critical for neuronal maturation and function. Indeed, mouse models of RTT demonstrate smaller brain size without detectable cell loss, reduction in number and length of dendrites, and a reduction in dendritic spine numbers^9,10^. These lines of evidence demonstrate the importance of MeCP2 in the proper regulation of synaptic development and function.

At the molecular level, a consensus of MeCP2 function and how its dysfunction leads to disease is currently lacking. MeCP2 contains five protein domains: N-terminal domain (NTD), methyl-CpG-binding domain (MBD), intervening domain (ID), transcriptional repression domain (TRD), and the C-terminal domain (CTD). Through the MBD, MeCP2 binds to methylated and hydroxy-methylated cytosines on genomic DNA^11,12^, and recruits co-repressors such as the Sin3A and NCoR complexes through its TRD^13,14^. In addition, an early study has shown that the TRD of MeCP2 represses transcription *in vitro*^15^. Based on these results, MeCP2 has traditionally been viewed as a repressor of transcription. However, several subsequent studies have suggested that MeCP2 may also play a role in activation of gene expression. First, transcriptomic studies in the hypothalamus of *Mecp2* null mice have shown that most of the differentially expressed genes are downregulated while some are upregulated, and those same genes are inversely altered in the overexpression mouse model^16^. This bi-directional inverse gene expression changes in mouse models of loss and gain of MeCP2 have been reproduced in followup studies^17,18^. Second, our lab has previously discovered that TCF20, which is associated with transcriptional activation, interacts with MeCP2^19^. Last, more recently, MeCP2 has been shown to bind to hypomethylated promoter-proximal regions and globally recruit RNA polymerase II (RNA pol II)^20^.

We rationalized that using an unbiased approach to identify the repertoire of functional interactors of MeCP2 would enable us to gain a deeper understanding of the function of MeCP2 at the molecular level. *Drosophila* lacks an orthologous gene to *MECP2* and shows minute levels of methylation on their genomic DNA^21^, yet the ectopic overexpression of human *MECP2* (h*MECP2*) in *Drosophila* can recapitulate its key functional characteristics. For example, overexpressed h*MECP2* associates with *Drosophila* chromatin, genetically interacts with *Drosophila* orthologs of known interacting partners including components of the Sin3A, NCoR, and SWI/SNF complexes, and can be phosphorylated at serine 423 as in mammals^22^. These lines of evidence suggest that the biology of MeCP2 can be interrogated in *Drosophila*, giving us a unique opportunity to use this genetic tool to find genetic interactors of *MECP2*.

Using the h*MECP2* gain-of-function *Drosophila* model, we knocked down chromatin-associated genes to determine if any could modify the h*MECP2*-induced rough eye phenotype. From this screen, we validated known interactors of MeCP2, but also discovered that knockdown of components of the super elongation complex (SEC) suppress the h*MECP2*-induced rough-eye phenotype. The SEC is a transcriptional activator complex composed of AFF1/AFF4, AF9/ENL, ELL1/ELL2, EAF1/EAF2, and P-TEFb which phosphorylates serine-2 of the carboxy-terminal domain (CTD) of promoter-proximally paused RNA pol II. Upon an external stimulus such as heat shock, the SEC releases RNA pol II into the genebody^23–25^. We find that MeCP2 directly interacts with AFF4, the scaffold of the SEC, and that the TRD is essential for this interaction. Furthermore, we find that MeCP2 facilitates the recruitment of AFF4 on a set of highly expressed genes associated with the regulation of synaptic function in the mouse brain. Coincidentally, RNA pol II binding was significantly depleted on the genebody of these genes in the cortex of *Mecp2* null mice, resulting in their downregulated expression. Collectively, these data raise the possibility that the interaction between MeCP2 and the SEC may facilitate transcription to regulate synaptic function.

## RESULTS

### Knockdown of components of the Drosophila SEC suppress the hMECP2-induced rough-eye phenotype

To find chromatin-associated genes whose knockdown could modify the h*MECP2*-induced rough eye, we screened through a total of 301 RNA interference (RNAi) lines that target 239 genes in *Drosophila* (**Fig. 1A**). From this screen, we identified 16 suppressor and 8 enhancer genes (**Fig. 1B**). Consistent with previous results^22^, we found that knockdown of previously known interactors modified the h*MECP2*-induced rough eye phenotype. Knockdown of components of the NCoR and SWI/SNF complexes mitigated, while knockdown of components of the Sin3A complex aggravated the h*MECP2*-induced rough eye phenotype (**Figs. 1C** and **S1A)**. Besides the known interactors, we also identified components of the SEC (*ear*, *Su(Tpl)*, *lilli*, and *Eaf*), *CG10565*, *Fbxl4*, Ada2a-containing (ATAC) complex (*D12*, *Ada2a*, and *Atac1*), RUVBL complex (*pont* and *rept*), *Taf10*, *ftz-f1*, and *Su(var)2-10* as new genetic interactors of h*MECP2* (**Fig. 1C**). Among the candidates, the SEC caught our attention for two reasons. First, individual knock down of all SEC components strongly suppressed the h*MECP2*-induced rough eye phenotype, suggesting there might be a strong genetic interaction between *MECP2* and the SEC. Second, since the SEC facilitates transcription, we hypothesized that exploring the functional interaction between MeCP2 and the SEC could reveal a previously unknown mechanism underlying MeCP2-mediated transcriptional regulation. For these reasons, we decided to focus on the interaction between MeCP2 and the SEC going forward.

**Figure 1.**
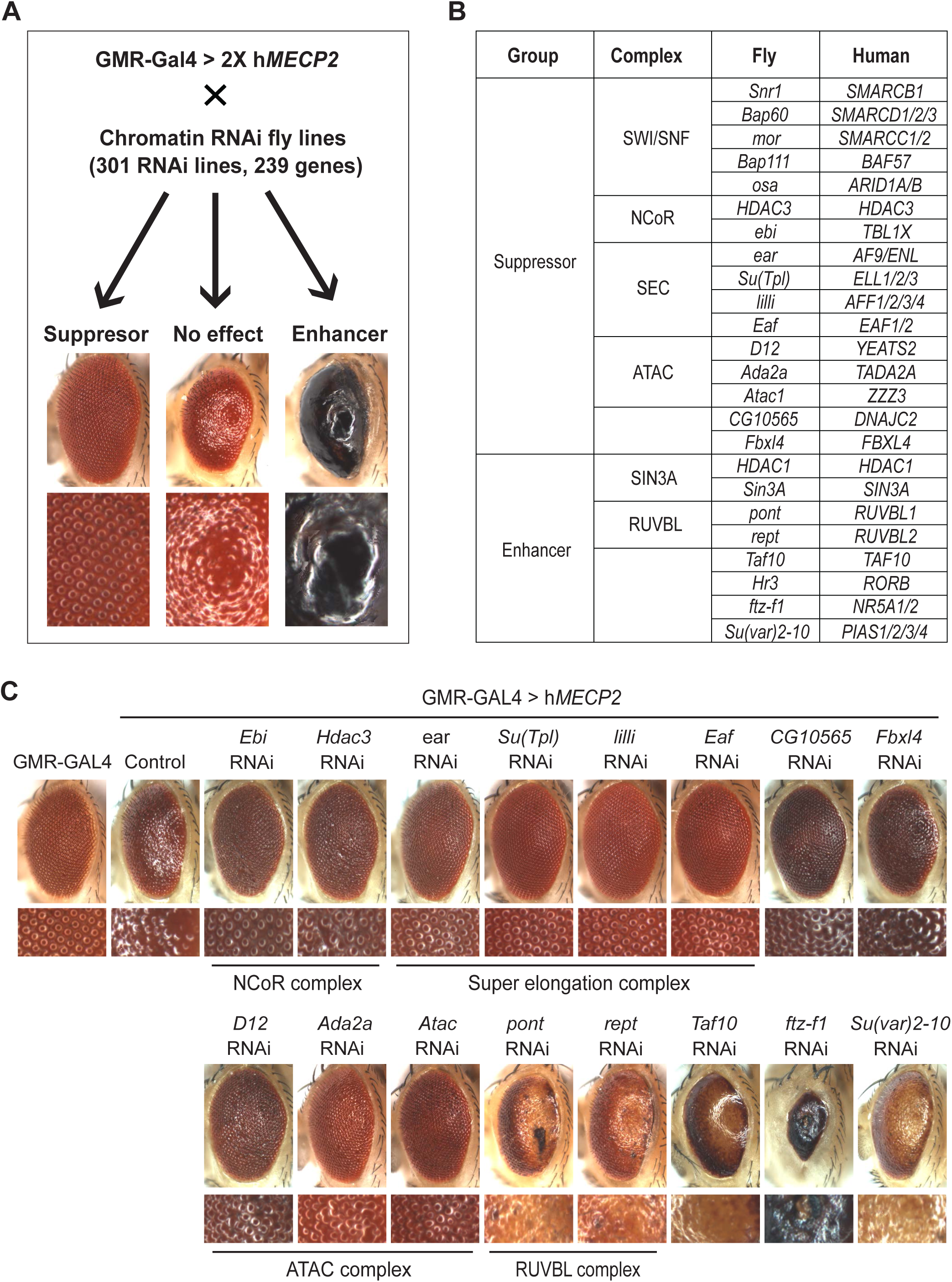
*MECP2* genetic modifier screen in *Drosophila*. (A) Schematic to find genetic modifiers of h*MECP2* in *Drosophila*. Images in bottom row represent magnifications of the respective images on the top. Images under “No effect” represent a control fly overexpressing two copies of h*MECP2*. (B) Table showing *Drosophila* candidate modifier genes and their human orthologues, knockdown of which either mitigates (suppressor) or aggravates (enhancer) the h*MECP2*-induced rough eye phenotype. (C) Light microscope images of eyes with no overexpression (GMR-GAL4) and overexpression of h*MECP2* (control) at 28.5°C. RNAi-mediated knockdown of NCoR complex components, SEC components, *CG10565*, *Fbxl4*, and ATAC complex components suppress, while knockdown of RUVBL complex components, *Taf10*, *ftz-f1*, and *Su(var)2-10* enhance the rough-eye phenotype. Image of *Hr3* knockdown is not shown as it caused lethality.

Before proceeding, we sought to determine if knockdown of components of the *Drosophila* SEC could affect the GAL4-UAS system and potentially result in an artificial rescue phenotype. In flies expressing GFP under the GMR-GAL4 driver, we knocked down SEC components via RNAi and measured the GFP protein levels in the head using Western blot. Knockdown of *Drosophila* SEC components using multiple RNAi resulted in the following GFP levels compared to the control (VK37): *lilli* = 17∼45%; *Su(Tpl)* = 66∼78%; *ear* = 93∼120%; and *Eaf* = 98%, while suppressing the h*MECP2*-induced rough eye phenotype (**Figs. S1B-S1C**). To exclude the possibility that knockdown of *lilli* suppressed the rough eye phenotype due to a down-regulation of GAL4-UAS activity, we tested the effect of a third RNAi line against *lilli* (*lilli* RNAi #3) on the h*MECP2*-induced rough eye phenotype and GAL4-UAS activity. We observed that *lilli* RNAi #3 decreased GFP levels by only ∼25% while strongly suppressing the rough eye phenotype (**Figs. S1D-S1E**). Collectively, these results suggest that an alteration in the genetic interaction between *MECP2* and the SEC is responsible for suppressing the h*MECP2*-induced rough eye phenotype.

### MeCP2 physically interacts with the SEC and RNA pol II

To determine if MeCP2 can physically associate with the SEC, we co-expressed C-terminal YFP-tagged MeCP2 (MeCP2-cYFP) with N-terminal YFP-tagged SEC components (AFF4-nYFP, ENL-nYFP, and EAF1-nYFP) in HEK293T cells and performed bimolecular fluorescence complementation (BiFC)^26^. Co-expression of MeCP2-cYFP with Ppp6c-nYFP (negative control), a protein that is not known to interact with MeCP2, did not produce YFP-positive cells when analyzed by fluorescence-activated cell sorting (FACS, **Fig. S2A**). However, co-expression of MeCP2-cYFP with TBL1X-nYFP (positive control), a component of the NCoR complex known to directly bind to MeCP2, resulted in the presence of YFP-positive cells, thereby demonstrating the specificity of our assay. When MeCP2-cYFP was co-expressed with nYFP-tagged SEC components, YFP-positive cells were detected, suggesting a physical interaction between MeCP2 and the SEC. Confocal microscopy images of the YFP signal further confirmed that MeCP2 interacts with components of the SEC within the nucleus (**Fig. S2B**).

Next, to determine if MeCP2 and the SEC interact in a physiological context, we immunoprecipitated (IPed) endogenous MeCP2 from HEK293T cell lysates and then probed for the co-immunoprecipitation (co-IP) of endogenous SEC components. We observed that AFF4, AF9, ENL (AF9 paralog), and ELL2 co-IPed with MeCP2 (**Fig. 2A**). Since the SEC phosphorylates RNA pol II for productive elongation, we rationalized that MeCP2 might also associate with RNA pol II. Indeed, we observed that both total RNA pol II and its elongating form (pSer2 RNA pol II) co-IPed with MeCP2.

**Figure 2.**
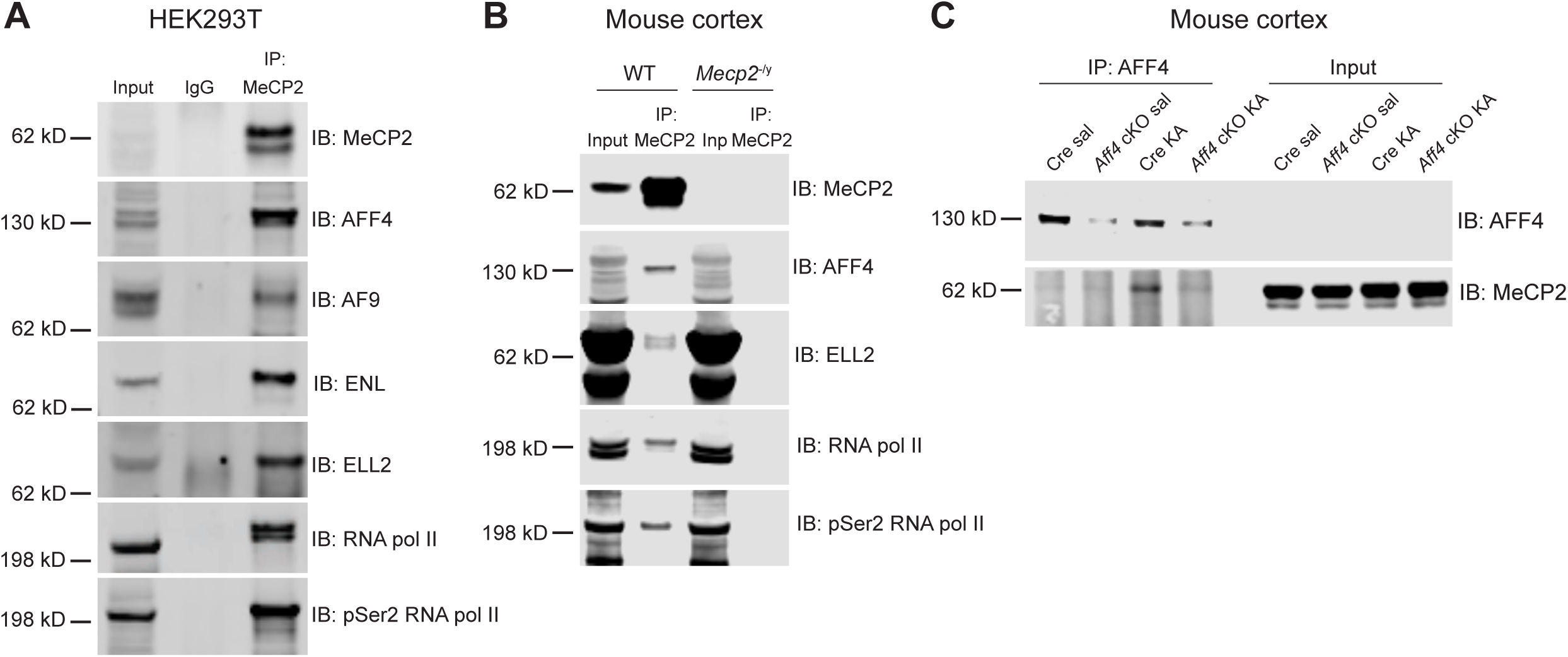
MeCP2 physically interacts with the SEC. (A) Endogenous MeCP2 interacts with endogenous SEC components (AFF4, AF9, ENL, and ELL2) and RNA pol II in HEK293T cells. Normal mouse IgG was used as a negative control. (B) Endogenous MeCP2 interacts with SEC components (AFF4 and ELL2) and RNA pol II in the cortex of WT mouse at 7-weeks of age. (C) Reverse IP of endogenous AFF4 from cortical lysates of Camk2a-Cre/+ control (Cre) and Camk2a-Cre>*Aff4*^fl/fl^ (*Aff4* cKO) mice at 7-weeks of age. Mice were injected with either saline (baseline) or kainic acid (KA).

To confirm if MeCP2 interacts with the SEC *in vivo*, we IPed MeCP2 from the cortex of wildtype (WT) and *Mecp2* null mice and probed for the co-IP of SEC components. We observed that AFF4 and ELL2 coIPed with MeCP2 in WT, but not in *Mecp2* null mice (**Fig. 2B**). To further validate this interaction, we reversely IPed for AFF4 and probed for the co-IP of MeCP2. However, we were only able to detect a faint MeCP2 co-IP band (**Fig. S2C**). We reasoned that since the SEC functions in a stimulus-dependent context^23–25^, the interaction between SEC and MeCP2 might be more pronounced after a stimulation. To this end, we stimulated neurons in the cortex of Camk2a-Cre (Cre control) and *Aff4* conditional knockout mice (cKO; Camk2a-Cre>*Aff4*^fl/fl^) through the injection of kainic acid (KA), a glutamate receptor agonist that depolarizes neurons. Interestingly, we observed a stronger co-IP of MeCP2 in KA-injected Cre control mice (**Fig. 2C**). This suggests that neuronal activity can enhance the interaction between MeCP2 and the SEC.

While the SEC utilizes AFF4 as its scaffold, two additional SEC-like complexes, SEC-like 2 (SEC-L2) and SEC-like 3 (SEC-L3) exist. These complexes utilize AFF2 and AFF3 paralogs as their scaffolds, respectively^27^. To determine if MeCP2 also interacts with SEC-L2 and SEC-L3, we IPed MeCP2 from cortical lysates of WT mice and probed for the co-IP of AFF2 and AFF3. However, we did not observe an interaction between these paralogs and MeCP2 (**Fig. S2D**), suggesting that MeCP2’s interaction is specific to the AFF4-associated SEC in the mouse cortex.

### The TRD is essential for MeCP2’s direct binding to AFF4

To determine which component of the SEC mediates the interaction with MeCP2, we knocked down three different components of the SEC (*AFF4*, *ENL*, and *ELL2*) in HEK293T cells and tested for the interaction between MeCP2 and the rest of the SEC components. Interestingly, knockdown of *AFF4* significantly reduced the co-IP between other components of the SEC (ENL and ELL2) and MeCP2 (**Fig. 3A**). In contrast, neither knocking down *ENL* nor *ELL2* affected MeCP2’s interaction with other SEC components, suggesting that AFF4 primarily mediates the interaction between MeCP2 and the SEC.

**Figure 3.**
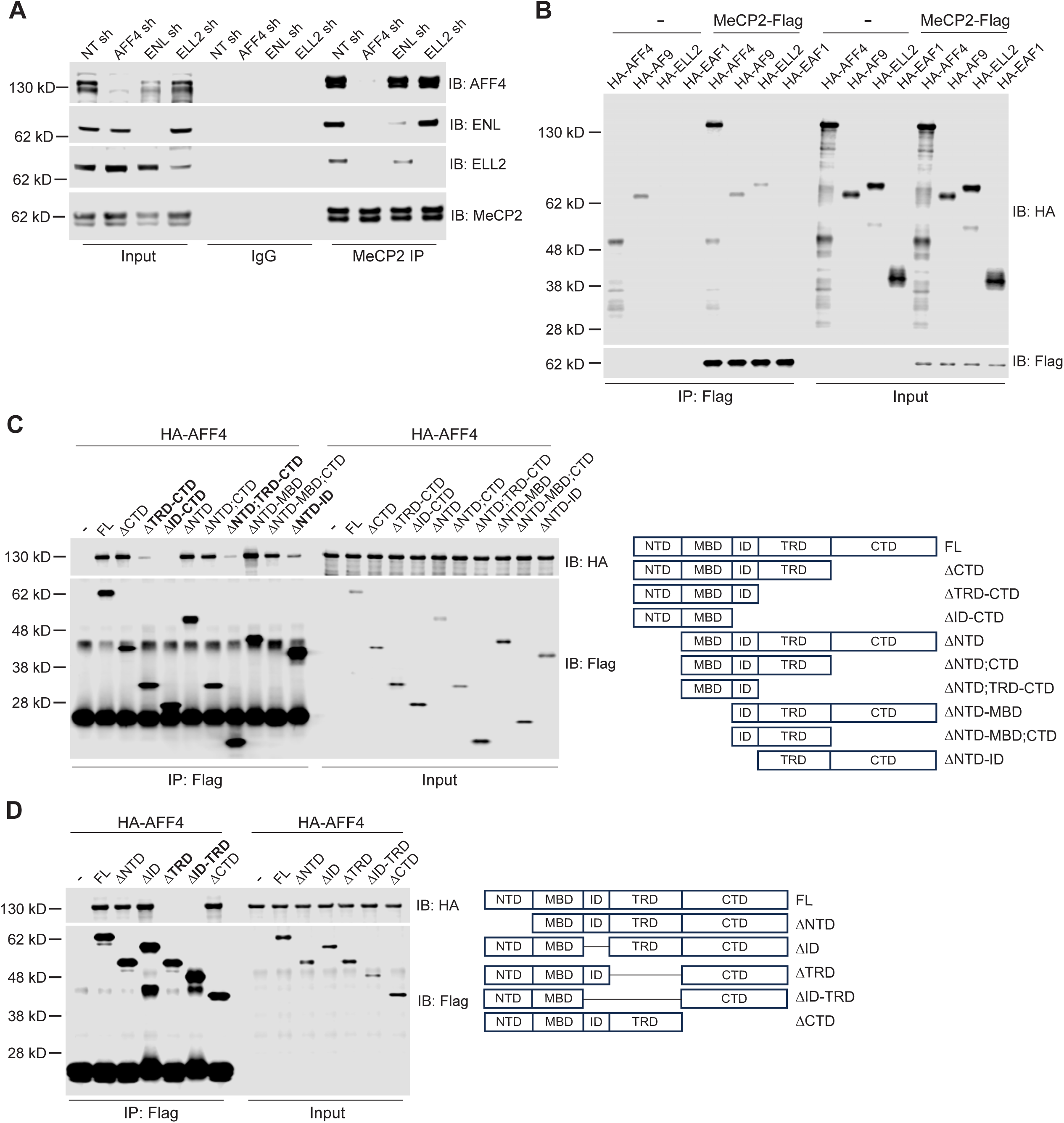
The TRD is required for MeCP2’s direct binding to AFF4. (A) shRNA-mediated knockdown of *AFF4* in HEK293T cells abolishes the interaction between MeCP2 and the remaining components of the SEC (ENL and ELL2). In contrast, knockdown of ENL or ELL2 does not affect the interaction between MeCP2 and the remaining components of the SEC. NT sh is a non-targeting short-hairpin negative control. (B) *In vitro* binding assay between recombinant FLAG-tagged MeCP2 and HA-tagged SEC components. (C) *In vitro* binding assay between FLAG-tagged MeCP2 fragments and HA-tagged AFF4. The domains critical for interaction are highlighted in bold. Diagram on the right shows a map of the MeCP2 fragments that were tested. NTD = N-terminal domain; MBD = methyl-CpG-binding domain; ID = intervening domain; TRD = transcriptional repression domain; CTD = C-terminal domain. (D) *In vitro* binding assay between AFF4 and MeCP2 with deletions in the NTD, ID, TRD, ID and TRD, and CTD. The domains critical for interaction are in bold.

In parallel, we also examined the direct binding between MeCP2 and SEC components *in vitro*. Using an *in vitro*-coupled transcription and translation (*in vitro* TnT) system, we generated FLAG-tagged MeCP2 (MeCP2-FLAG) and HA-tagged SEC components (HA-AFF4, HA-AF9, HA-ELL2, and HA-EAF1). Subsequently, MeCP2-FLAG was incubated with each of the recombinant SEC components followed by IP using FLAG antibody-conjugated beads. We observed that HA-AFF4 did not come down in the absence of MeCP2-FLAG and only came down when MeCP2-FLAG was present (**Fig. 3B**). Further, we observed that HA-AF9 came down even in the absence of MeCP2-FLAG, indicating that FLAG-conjugated beads non-specifically bound to AF9. However, in the presence of MeCP2-FLAG, HA-AF9 did not come down more, suggesting that MeCP2 does not bind to AF9. We noticed a small amount of HA-ELL2 came down with MeCP2-FLAG, suggesting that MeCP2 weakly interacts with ELL2. Finally, HA-EAF1 did not come down either in the absence or presence of MeCP2-FLAG, indicating that EAF1 does not interact with MeCP2. Collectively, these results suggest that MeCP2 preferentially and directly binds to AFF4.

Next, to identify the protein domain in MeCP2 that is required for its interaction with AFF4, we generated FLAG-tagged MeCP2 fragments with various domain deletions using the *in vitro* TnT system and incubated the recombinant fragments individually with HA-AFF4. We found that deleting the NTD, CTD, and MBD did not affect MeCP2’s interaction with AFF4 (**Fig. 3C**). Interestingly, we discovered that deletion of the TRD greatly reduced MeCP2’s binding to AFF4. In addition, we noticed that deletion of the ID with other domains mildly impacted the binding. However, when we deleted only the ID, it did not affect MeCP2’s binding to AFF4 (**Fig. 3D**). In contrast, deletion of only the TRD greatly reduced the binding. Collectively, these results indicate that the TRD of MeCP2 is essential for its interaction with AFF4.

### MeCP2 facilitates the binding of AFF4 on a subset of genes while promoting the global binding of RNA pol II

Given that MeCP2 physically interacts with the SEC, we hypothesized that MeCP2 recruits the SEC to facilitate RNA pol II binding. To test this hypothesis, we decided to assess whether the loss of MeCP2 can affect the genome-wide binding of AFF4 and RNA pol II in the mouse brain using chromatin immunoprecipitation sequencing (ChIP-seq). Before proceeding, we first tested whether our antibody could specifically detect AFF4 in the mouse brain. Since AFF4 binding is induced on genes that respond to stimulus^23–25^, we decided to evaluate the binding of AFF4 in the KA-activated brain. To this end, we injected either saline or KA into Cre control (Camk2a-Cre/+) and *Aff4* cKO (Camk2a-Cre>*Aff4*^fl/fl^ ) mice and performed AFF4 ChIP-seq on the cortical lysates. We observed that a short treatment (45 minutes) of KA in the Cre control mice induced AFF4 binding on a small cluster of genes, including activity-dependent genes such as *Homer1* and *Egr3* (**Figs. S3A-S3B**). *Aff4* cKO mice failed to show this induction of binding on a subset of the gene cluster. It is likely that we could not observe an abolishment of the KA-induced binding of AFF4 on all the genes in the cluster because AFF4 binding was also induced on genes that are expressed in cell types besides the Camk2a-positive excitatory neurons, including inhibitory neurons, pericytes, and endothelial cells (**Fig. S3C**). Nevertheless, the fact that we see a significant suppression of the KA-induced binding of AFF4 on activity-dependent genes demonstrates the validity of our antibody.

Comparing the genome-wide binding of AFF4 in the cortex of WT and *Mecp2* null mice, we observed a reduction in binding of AFF4 on a subset of genes (**Figs. 4A-4B** and **Fig. S4A**). To ensure this reduction in binding was not due to a decrease in protein levels of AFF4, we measured AFF4 levels in the cortex of WT and *Mecp2* null mice via Western blot. We observed that AFF4 protein levels were comparable between WT and *Mecp2* null mice (**Fig. S4B**), suggesting that the decreased binding of AFF4 is not due to a decrease in protein levels. Additionally, when we compared the genome-wide binding of RNA pol II, surprisingly, we found a global reduction in *Mecp2* null mice (**Figs. 4C-4D** and **Fig. S4C**). Collectively, these results suggest that MeCP2 globally supports the binding of RNA pol II and the interaction between MeCP2 and the SEC contributes to support this binding on a subset of genes.

**Figure 4.**
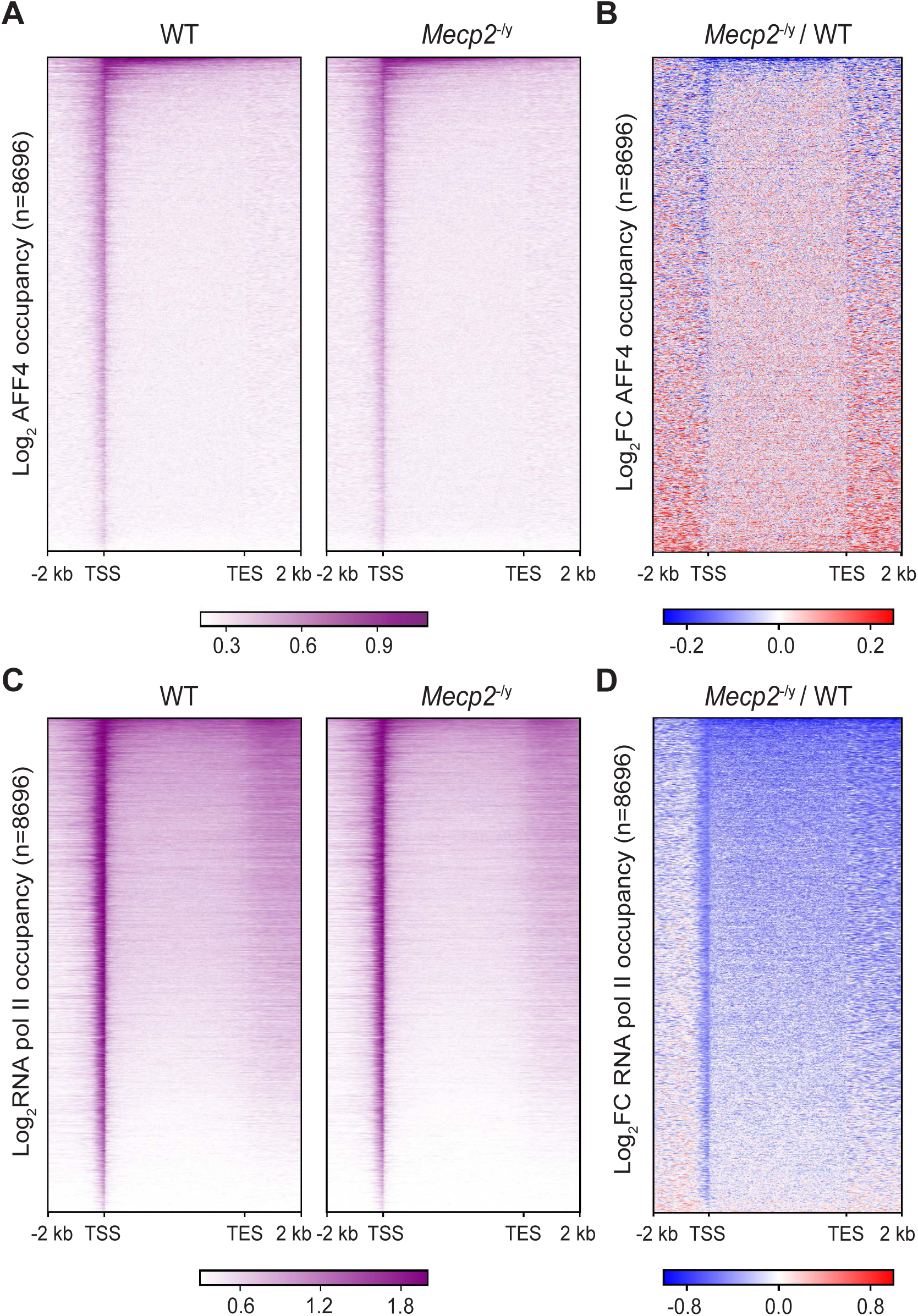
MeCP2 facilitates the binding of AFF4 on a subset of genes while globally promoting the binding of RNA pol II. (A) Heatmap of log occupancy of AFF4 in the cortex of WT and *Mecp2* null mice. (B) Heatmap of log fold change of AFF4 occupancy in *Mecp2* null mouse compared to WT mouse. (C) Heatmap of log occupancy of RNA pol II in the cortex of WT and *Mecp2* null mice. (D) Heatmap of log fold change of RNA pol II occupancy in *Mecp2* null mouse compared to WT mouse. 8696 RNA pol II-bound genes are represented in all heatmaps.

### MeCP2 supports the expression of SEC-regulated genes

To dissect the interplay between MeCP2, SEC, and RNA pol II, we clustered the genes based on the change of AFF4 and RNA pol II binding in *Mecp2* null mice. This generated three independent clusters: clusters I, II, and III (**Fig. 5A** and **Fig. S5A**). Cluster I (n = 2104 genes) displayed a mixture of genes that exhibited both increased and decreased binding of AFF4 upstream of the transcription start site (TSS) and downstream of the transcription end site (TES). For RNA pol II, genes in this cluster showed a decrease in binding on the TSS. Cluster II (n = 6263 genes) displayed a similar mixture of genes with both increased and decreased binding of AFF4 upstream of the TSS and downstream of the TES. However, RNA pol II binding was decreased not only on the TSS, but also across the genebody and beyond the TES. Last, although cluster III was comprised of only 329 genes, AFF4 binding was decreased on the TSS and across the genebody and demonstrated the strongest reduction in the binding of RNA pol II across the genebody. To determine if the change in AFF4 and RNA pol II binding in *Mecp2* null animals were correlated to each other, we plotted the genebody change in AFF4 versus RNA pol II binding across the three clusters. Although Spearman’s correlation analysis did not reveal a meaningful correlation in any of the clusters (cluster I: ρ=0.14, p=1.5e-10; cluster II: ρ=0.09, p=8.2e-10; cluster III: ρ=0.09, p=0.1), we observed a trend in which the proportion of genes that have a decrease in both AFF4 and RNA pol II binding (quadrant 3) sequentially increased from cluster I to cluster III (cluster I = 36%; cluster II = 55%; cluster III = 84%, **Fig. 5B**). This observation raises the possibility that the decreased binding of AFF4 due to loss of MeCP2 may contribute to the decreased binding of RNA pol II in the genebody of cluster III genes.

**Figure 5.**
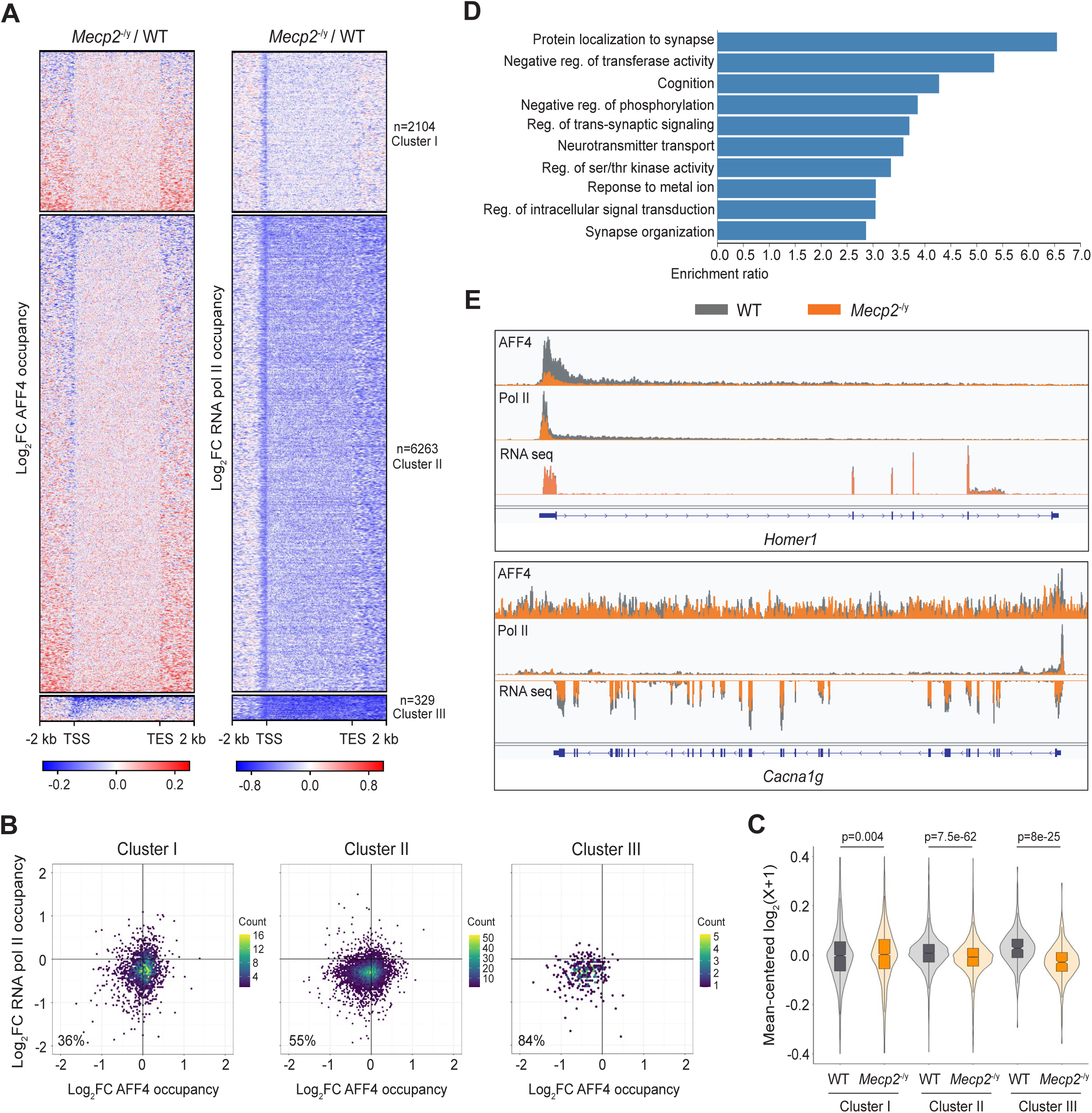
MeCP2 supports the expression of SEC-regulated genes. (A) Heatmap showing the log fold change of AFF4 and RNA pol II binding in the *Mecp2* null cortex based on hierarchical clustering. (B) Two-dimensional plot showing the change in AFF4 vs. RNA pol II binding in *Mecp2* null animals across each cluster. Color scale indicates the gene count. Data shown is the average of three biological replicates. Spearman’s correlation values: cluster I ρ=0.14, p=1.5e-10; cluster II ρ=0.09, p=8.2e-10; cluster III ρ=0.09, p=0.1. (C) Violin plot showing the average expression value of all genes in each cluster relative to WT cluster I. Data shown is the average of six biological replicates. Statistical significance was assessed using the Wilcoxon Rank Sum Test. (D) Gene ontology analysis of cluster III genes. All terms are FDR < 0.05. (E) Track examples of AFF4 and RNA pol II binding and RNA expression levels in WT (grey) and *Mecp2* null (orange) mouse.

To determine if the change in binding of AFF4 correlates with the change in gene expression, we also performed RNA-seq on the cortex of WT and *Mecp2* null mice. Principal component analysis revealed that the WT samples segregated from the *Mecp2* null samples (**Fig. S5B**), demonstrating the validity of our RNA-seq results. When we compared the average expression of genes in each cluster, we found that cluster I had a slight increase while clusters II and III were decreased in *Mecp2* null mice, with cluster III showing the greatest magnitude of change (**Fig. 5C**). When we plotted the change in AFF4 binding versus the change in RNA expression in *Mecp2* null animals across the three clusters, again we did not detect a meaningful correlation (cluster I: ρ=0.02, p=0.36; cluster II: ρ=0.047, p=0.00023; cluster III: ρ=0.15, p=0.0076), but again observed that cluster III had the highest proportion of genes (61%) that showed a decrease in both AFF4 binding and RNA expression (**Fig. S5C**). Interestingly, our analysis of the binding levels of AFF4 and RNA pol II along with RNA expression levels across the three clusters in WT revealed that cluster III genes not only had the highest binding levels of AFF4 and RNA pol II, but also had the highest RNA expression level (**Figs. S5D-S5E**). This observation indicates that cluster III is comprised of the most highly expressed genes in the mouse cortex, which are under the regulation of the SEC. Gene ontology analysis of cluster III revealed an enrichment of genes whose protein products are involved in the regulation of synaptic function such as localization of protein to synapse, trans-synaptic signaling, and neurotransmitter transport (**Figs. 5D-5E**). Collectively, these data suggest that MeCP2 facilitates the recruitment of the SEC to regulate synaptic function in the mouse brain.

## DISCUSSION

Through an unbiased genetic modifier screen in *Drosophila*, we have not only confirmed previously known interactors of MeCP2 such as components of the Sin3A, NCoR, and SWI/SNF complexes, but also discovered several new interactors. This underscores the reliability and power of our screening approach. From this screen, we identified the SEC as a new interactor of MeCP2. MeCP2 physically interacts with the SEC by directly binding to AFF4 and facilitates its recruitment to a subset of genes involved in regulating synaptic function. We also found that a large portion of these genes exhibit reduced RNA pol II binding, particularly on the genebody, and also exhibits down-regulated expression. Collectively, these findings shed light on a previously unknown mechanism by which MeCP2 recruits the SEC to regulate transcription.

Previous studies have demonstrated that neuronal activation can induce phosphorylation of MeCP2 at multiple amino acid residues in the rodent brain^28,29^. Among these, activity-dependent phosphorylation on threonine 308 has been shown to dissociate the interaction between MeCP2 and the NCoR complex^30^. To the best of our knowledge, our study is the first to demonstrate that MeCP2 can acquire a new interactor upon neuronal activation. We have found that the association of AFF4 with MeCP2 increases upon activation of excitatory neurons. Additionally, our result demonstrates that KA-induced activation of the mouse cortex increases AFF4 binding on a specific set of activity-dependent genes. Taken together, these observations raise the possibility that the interaction between MeCP2 and the SEC may serve to regulate activity-dependent genes when neurons become excited to potentially modulate synaptic plasticity. Future studies that dissect the functional interaction between MeCP2 and the SEC in a neuronal activity context would be interesting. More broadly, the ability of MeCP2 to acquire a new interactor upon neuronal activation suggests that MeCP2 may reorganize its protein interactors based on the activity state of neurons. To gain a deeper understanding of the molecular mechanisms underlying MeCP2-dependent transcription in neurons, characterizing the interactors of MeCP2 in both baseline and activity states would be very useful. At the protein level, we find that MeCP2 directly binds to AFF4 through the TRD. Notably, the TRD in MeCP2 is known to bind to other interactors, such as the NCoR co-repressor complex^13,14^. This raises the question of whether co-repressor and co-activator interactors of MeCP2 occupy different regions within the TRD or if they compete for the same region. Further detailed mapping of the specific region within the TRD that mediates the interaction between MeCP2 and AFF4 will be critical to address this important question.

There has been a long-standing controversy in the field regarding the mechanism by which MeCP2 regulates transcription. MeCP2 has been proposed to function as a roadblock on the genebody, impeding RNA pol II progression and thereby regulating its elongation speed^31,32^. However, the study by Boxer *et al*. which measured the rate of RNA pol II elongation in the forebrain of wildtype and *Mecp2* null mice, showed that MeCP2 does not impact the speed at which RNA pol II travels along the genebody^33^. Instead, MeCP2 was proposed to repress the transcriptional initiation of highly methylated long genes. Recently, Yi *et al*. reported that MeCP2 occupies CpG-rich promoter-proximal regions and globally recruits RNA pol II in human ipsc-derived neurons^20^. In agreement with Yi *et al*.’s finding, we provide here *in vivo* evidence that MeCP2 globally supports the binding of RNA pol II. Out of 8,696 RNA pol II-bound genes that were analyzed, approximately 90% of them showed decreased RNA pol II binding in the *Mecp2* null cortex. Interestingly, we find that loss of MeCP2 affects RNA pol II binding in two distinct ways. In cluster I, which primarily consists of paused genes with minimal expression, RNA pol II binding is depleted mostly on the TSS. Conversely, in clusters II and III, which consist of actively transcribing genes, RNA pol II binding is depleted both on the TSS and across the genebody. This observation leads to the hypothesis that MeCP2 influences transcription at both the initiation and elongation stages by recruiting different protein partners. While transcriptional initiation factors that functionally interact with MeCP2 have yet to be identified, we propose that MeCP2 facilitates transcriptional elongation by recruiting the SEC to genes. Future studies assessing the transcriptional dynamics in MeCP2-depleted neurons using nascent transcription assays will be critical to understand, with greater resolution, how MeCP2 regulates transcription.

In summary, our study has utilized an unbiased genetic modifier screen in *Drosophila* to uncover new genetic interactors of *MECP2*. These findings reveal that the molecular function of MeCP2 is complex, as it can interact with various proteins with diverse functions. This complexity, in turn, suggests that MeCP2 loss-of-function might disrupt interactions with multiple partners, which may all contribute to the pathogenesis of RTT. By establishing a network of genetic interactors of *MECP2*, we have laid the groundwork to further dissect the molecular function of MeCP2, which hopefully will allow us to gain a deeper understanding of the disease mechanisms underlying RTT.

## MATERIAL AND METHODS

### Drosophila genetic interactor screen

For the screen, we used transgenic UAS-h*MECP2* (2^nd^ and 3^rd^ chromosome) to construct GMR-Gal4, UAS-h*MECP2*/Cyo; UAS-h*MECP2*/TM6B flies. RNAi alleles were purchased from the Vienna *Drosophila* RNAi Center and the Bloomington Drosophila Stock Center repositories. Flies were crossed at 28.5°C and the external eye phenotype was imaged with a light microscope.

### Animals

All mice used in this study were maintained in a 12-hr light:dark cycle at 20-22°C and 30-70% humidity, with standard chow and water ad libitum. *Mecp2*^-/y^ mice, described previously^34^, were maintained on the 129S6/SvEvTac background. Constitutive *Aff4*^+/-^ mice (MMRRC; 041408-UCD) were maintained on the 129S6/SvEvTac background. *Aff4* flox mice, described previously^35^, were maintained on the C57BL/6J background and the Camk2a-Cre mouse was purchased from The Jackson Laboratory (JAX; 5359). For kainic acid injection, kainic acid (Tocris; 0222) was dissolved in 0.9% sodium chloride solution (Henry Schein; 1047098) and 25mg/kg was injected intraperitoneally. The Baylor College of Medicine Institutional Animal Care and Use Committee approved all research and animal care procedures.

### Cell culture

Human Embryonic Kidney 293T (HEK293T) cells were cultured with DMEM (VWR; 10-013-CV) supplemented with 10% heat-inactivated fetal bovine serum (FBS) and antibiotic-antimycotic (Thermo Fisher; 15240112). Cells were maintained at 37°C and 5% CO2.

### BiFC

Human *MECP2* (E2 isoform) cDNA was cloned into the pBiFC vector that tags the C-terminal half of YFP on the C-terminus. Human *AFF4*, *ENL*, and *EAF1* cDNAs were cloned into the pBiFC vector that tags the N-terminal half of YFP on the C-terminus. 0.5μg of pBiFC constructs expressing *MECP2* and SEC components were transfected into HEK293T cells using Lipofectamine 3000 (Thermo Fisher; L3000150) according to the manufacturer’s instruction. After 48 hours, cells were harvested for either FACS or confocal imaging.

Cells were washed once with PBS, then incubated with the Helix NP™ NIR dye (Biolegend; 425301), a nucleic acid stain used to discriminate live and dead cells. Cells were acquired and analyzed using a BD LSRFortessa flow cytometer (BD Biosciences). Unstained controls were included to set the baseline fluorescence and to determine background signal. A positive control sample expressing MeCP2-GFP was utilized to set the positive gate and a negative control sample expressing MeCP2-cYFP together with empty-nYFP was utilized to set the negative gate. FACSDiva software was used to collect the data, while FlowJo (v10.8) was used to perform analyses and generate relevant plots.

For confocal imaging, cells were washed once with PBS and then fixed with 4% formaldehyde. After fixation, cells were permeabilized with PBS containing 0.5% Triton-X and then stained with DAPI (Thermo Scientific; 62248). Coverslips with cells were mounted on slides with mounting medium (Thermo Fisher; P36961) and images were taken with a confocal microscope (Zeiss LSM710).

### Co-immunoprecipitation

For HEK293T cells, a confluent 15cm plate was lysed in 1mL of immunoprecipitation (IP) buffer (10 mM HEPES pH 7.9, 3 mM MgCl2, 5 mM KCl, 140 mM NaCl, 0.1mM EDTA, 0.5% NP-40) that was supplemented with protease and phosphatase inhibitors (Gendepot; P3100-020 and P3200-020, respectively) and 1:1,000 Pierce Universal Nuclease (Thermo Fisher; 88702). For cortical tissues, a single hemisphere was dounced in 1mL of IP buffer supplemented with protease and phosphatase inhibitors and nuclease. Lysates from cells and tissues were rotated on an end-over-end rotator for one hour at 4°C and then centrifuged at 16,000 rcf at 4°C for 20 minutes. The supernatant was transferred to a new tube and input was retrieved before mixing with Dynabead G (Thermo Fisher; 10004D, 20μl per IP for cells and 30 μl per IP for tissues) that were pre-conjugated with one of the following: 5μg of MeCP2 antibody (Sigma-Aldrich; M6818), 5μg AFF4 antibody (Proteintech; 14662-1-AP), and 5μg of normal mouse or rabbit IgG (Millipore; 12-371 and 12-370). The supernatant/Dynabead-antibody mixture was rotated overnight at 4°C. The following day, the beads were washed four times with 1ml of cold IP buffer and subsequently eluted in 25μl of 1.1X LDS sample buffer/reducing agent (Thermo Fisher; NP007 and NP009, respectively) at 95°C for 5 minutes with shaking.

### In vitro binding assay

Full-length MECP2 and its fragment cDNAs were cloned into the pT7CFE1 vector that tags FLAG on the C-terminus. SEC component cDNAs were cloned into the pT7CFE1 vector that tags HA on the N-terminus. The constructs were used for *in vitro* transcription and translation using the 1-Step Human High-Yield Mini IVT Kit (Thermo Fisher; 88891) according to the manufacturer’s instruction. Proteins to be tested for binding were incubated together in 250μl of IP buffer and incubated for one hour at 4°C on an end-over-end rotator. Before IP, 15μl were taken for input. 10μl of the FLAG-bead slurry (Sigma Aldrich; M8823-1ML) was reconstituted in 50μl of IP buffer per sample and added to the IP samples. After one hour of rotation at 4°C, beads were washed four times with IP buffer. Beads were eluted with 25μl of 1.1X LDS sample buffer/reducing agent at 95°C for 5 minutes with shaking.

### Western blot

Cell and tissue lysate samples were run on 4-12% Bis-Tris midi-gels (Invitrogen; WG1403BOX) and transferred to nitrocellulose membranes using wet transfer. Membranes were blocked for 30 minutes using the INTERCEPT (TBS) Blocking Buffer (Li-Cor; 927-60010) and then incubated with the following primary antibodies overnight at 4°C: AFF4 (Bethyl; A302-538A at 1:1,000 and Proteintech; 14662-1-AP at 1:5,000), AF9 (Genetex; GTX 102835 at 1:1000), ENL (Cell Signaling Technology; 14893 at 1:1000), ELL2 (Bethyl; A302-505A at 1:1000), total RNA pol II (Cell Signaling Technology; 14958S at 1:1000), pSer2 RNA pol II (Millipore; 04-1571 at 1:1000), MeCP2 (Cell Signaling Technology; 3456S at 1:1000), GFP (Abcam; ab13970 at 1:10,000), b-actin (Cell Signaling Technology; 8457S at 1:10,000), vinculin (Sigma Aldrich; V9131-.2ML at 1:20,000), FLAG (Sigma Aldrich; F3165-1MG at 1:4,000), and HA (Cell Signaling Technology; 3724S at 1:4,000). Membranes were washed three times with TBS-T, incubated with IRDye secondary antibodies (Li-COR; 926-68020, 926-32211, and 926-68028 at 1:10,000) and Alexa Flour 790-AffiniPure antibody (Jackson ImmunoResearch; 115-655-174 at 1:10,000) for one hour, and then washed three times with TBS-T. Blots were imaged on the Odyssey CLx (LiCoR).

### Chromatin immunoprecipitation sequencing

*Mecp2*^-/y^ (129S6/SvEvTac) female mice were bred to wildtype (FVB) males and the cortex was harvested at 7 weeks of age. Tissues were immediately snap-frozen in liquid nitrogen and stored at -80°C until use. One hemisphere of the cortex was used for each ChIP. Tissues were crosslinked with 1% formaldehyde in PBS for 10 minutes at room temperature and were quenched with 0.2M glycine for 5 minutes. Chromatin was sonicated with the Covaris E220 using the following conditions: 6 minutes, 5% duty cycle, 140 peak intensity power, and 200 cycles per burst for AFF4 and 6 minutes, 10% duty cycle, 140 peak intensity power, and 200 cycles per burst for RNA pol II. The following amounts of *Drosophila* spike-in chromatin (Active Motif; 53083) were added per ChIP: 35ng for AFF4 and 180ng for RNA pol II. 2μg of spike-in antibody (Active Motif; 61686) was added to each sample with one of the following antibodies for immunoprecipitations: 5μg of AFF4 antibody (Bosterbio; M03824) and 5μl of RNA pol II antibody (Cell Signaling Technology; 14958S). Immunoprecipitations were performed overnight with Dynabeads Protein G. Reverse crosslinking and protein degradation were performed at 65°C overnight. Immunoprecipitated DNA was purified using the QIAquick PCR purification kit (Qiagen; 28106). DNA libraries were prepared using the KAPA Hyperprep kit (Roche; KK8504). The cycle number for library amplification was empirically determined using SYBR Green I Nucleic Acid Gel Stain (Thermo Fisher; S7585) and real-time qPCR tracking (Biorad CFX96). Pooled libraries were sequenced on the NovaSeq 6000 (Illumina) using single-end 100 cycle or paired-end 50 cycle reads.

### Total RNA sequencing

*Mecp2*^+/-^ (129S6/SvEvTac) female mice were bred to wildtype (FVB) males and the cortex was harvested at 7 weeks of age. Tissues were immediately snap-frozen in liquid nitrogen and stored at -80°C until use. One cortical hemisphere was used to extract RNA using the RNeasy kit (Qiagen; 74106). Briefly, tissues were homogenized in buffer RLT using a tissue homogenizer (Cole Parmer). Homogenate was passed through a QIAshredder (Qiagen; 79656) and the flow through was processed with the RNeasy kit (Qiagen; 74104). Ribosomal RNA was depleted using the NEBNext rRNA Depletion Kit v2 (NEB; E7400L) and libraries were prepared using the NEBNext Ultra II Directional RNA Library Prep Kit for Illumina (NEB; E7770L). Pooled libraries were sequenced on the NovaSeq 6000 instrument using paired-end 51 cycle reads.

### Bioinformatic analysis

For ChIP-seq, reads were trimmed using Cutadapt v4.1 with parameters --nextseq-trim=30 --minimum-length=20, then aligned to the mm10 mouse genome assembly and dm6 *Drosophila* genome assembly for spike-in read mapping using Bowtie2 v2.2.6^36^ with parameters --sensitive --no-unal. Peaks were called using MACS2^37^ with parameters -q 0.05 -- keep-dup auto --nomodel, then annotated using HOMER v4.1.1. Reads from mouse fragments were normalized to the *Drosophila* spike-in reads. Protein coding gene detection and selection, TSS and gene-body region selection, and coverage mapping were performed using in-house R and perl scripts. We filtered genes based on the following criteria: 1) RNA pol II occupied genes, 2) longer than 2 kilobases (kb), and 3) neighboring genes that are at least 2kb apart. This resulted in a total of 8696 genes that were used for downstream analysis. Occupancy and log_2_ fold change heatmaps were generated using deepTools v.3.5.1^38^. For the hierarchical clustering analysis, we clustered based on two of three replicates in which the *Mecp2* null samples showed the greatest difference from WT samples.

For total RNA-seq, the output data were processed with bcl2fastq. Sequence quality was assessed using FastQC v 0.11.2, and quality trimming was done using Trimmomatic^39^. RNA-seq reads were aligned to the mm10 genome using STAR v.2.5.2^40^ and only uniquely mapped reads with a two-mismatch threshold were considered for downstream analysis, then quantified to the gene level using HTSeq^41^. Output bam files were converted into bigwig track files to display coverage throughout the genome (in RPM) using the GenomicRanges package^42^ as well as other standard Bioconductor R packages. Gene count tables were constructed following Ensembl gene annotations and used as input for RUVseq^43^ and differential gene expression analysis was performed using DESeq2^44^. Genes with adjusted p-values (Benjamini-Hochberg) of less than 0.05 were treated as differentially expressed.

## ACKNOWLEDGEMENTS

We thank Rebecca Meyer-Schuman, Ashley G. Anderson, Sameer S. Bajikar, and Larissa Nitschke for critical reading of the manuscript and Yaling Sun for all the genotyping. We would also like to thank Benjamin C. Howard for assisting with the next-generation sequencing work. This work was supported by Howard Hughes Medical Institute (H.Y.Z.), NINDS; R01NS057819 (H.Y.Z.), NICHD; P50HD103555 (BCM-IDDRC), and NCI; R50CA265372 (M.I.) and R35-CA197569 (A.S.).

## AUTHOR CONTRIBUTIONS

J.Y.S., W.K., A.S., and H.Y.Z. designed the experiments. J.Y.S., W.K., Y.A., Y.L., and L.P. performed the experiments. M.I. performed the bioinformatic analysis. J.Y.S., A.S., and H.Y.Z. interpreted the data. J.L.B. and L.W. provided the *Aff4* flox mouse line. J.Y.S. wrote the manuscript with I.A., J.B., A.S., and H.Y.Z. giving critical input. A.S. and H.Y.Z. supervised the research.

