## Supplementary figures and images for "MeCP2 Interacts with the Super Elongation Complex to Regulate Transcription"

### Supplemental Figures

#### Fig. S1 Sonn et al.

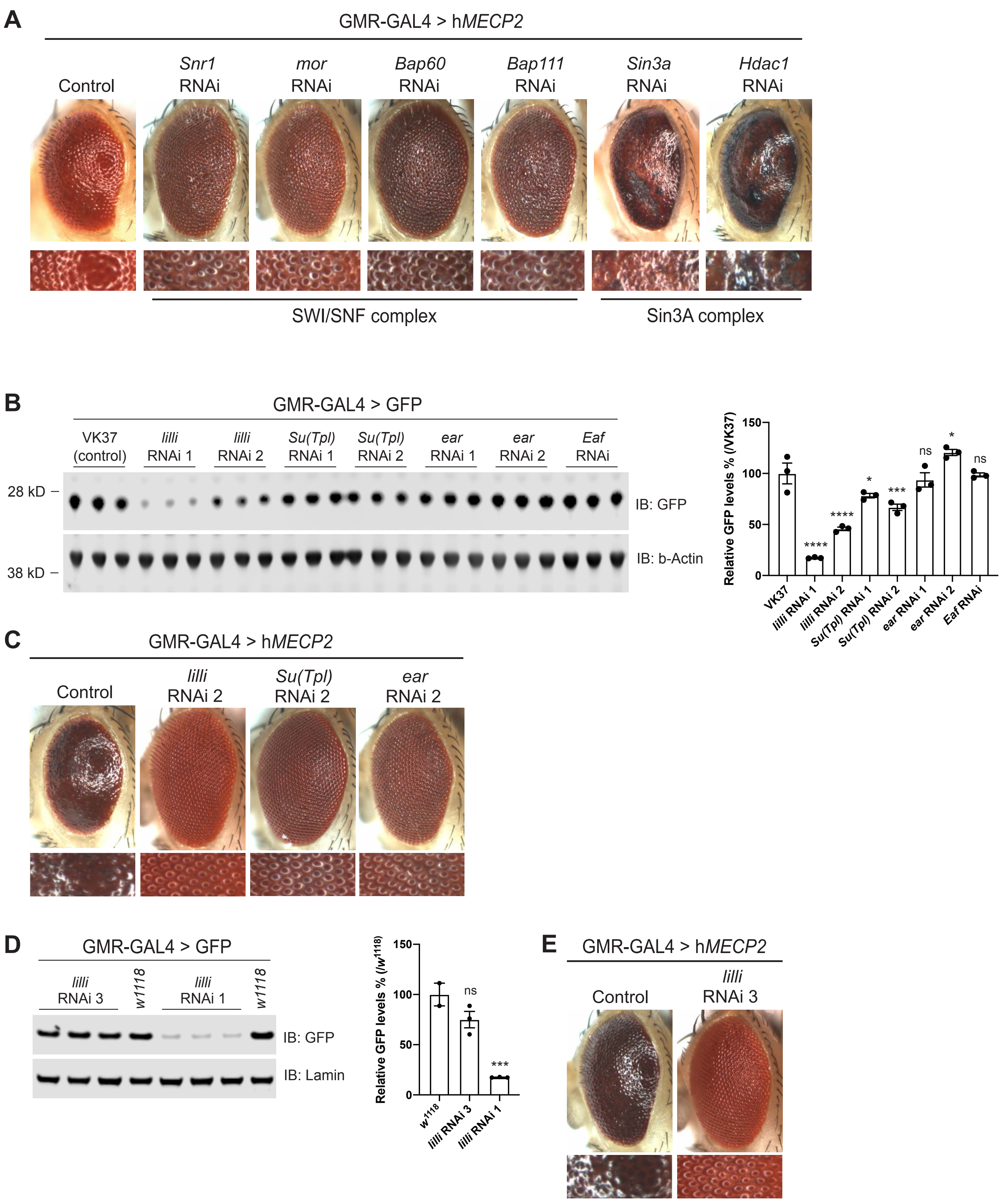

#### Fig. S2 Sonn et al.

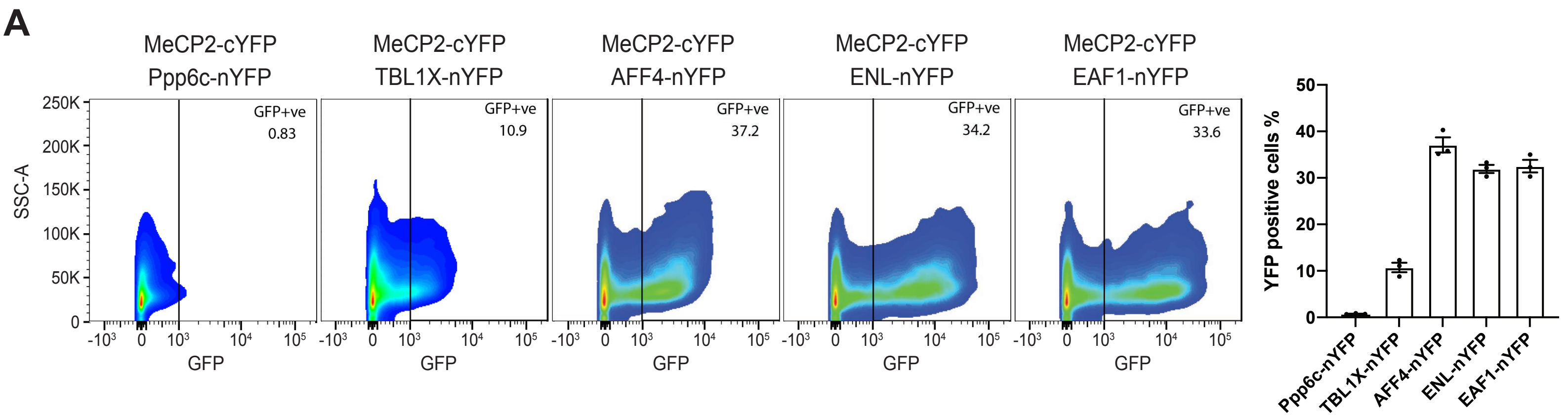


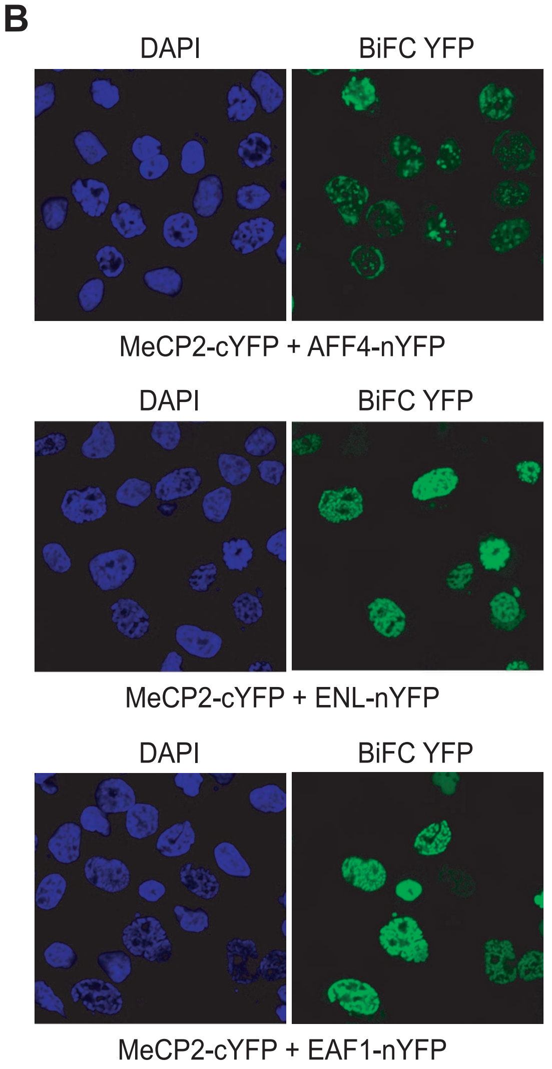


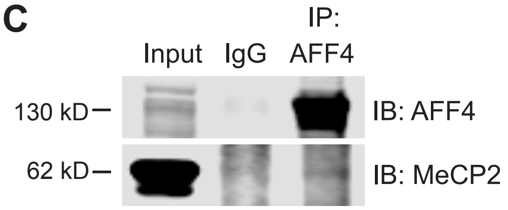


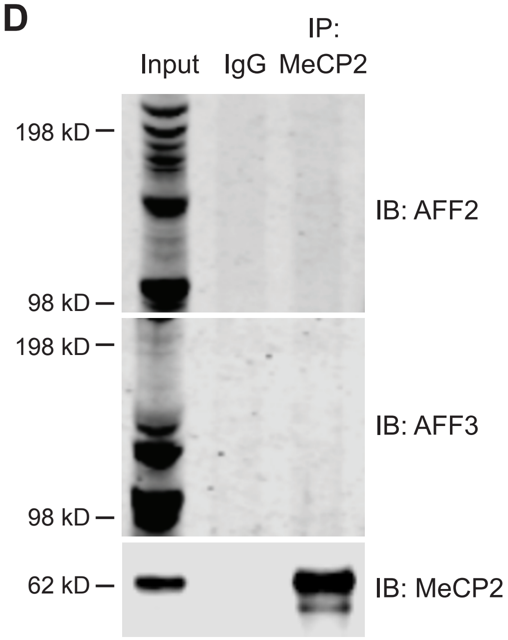

#### Fig. S3 Sonn et al.

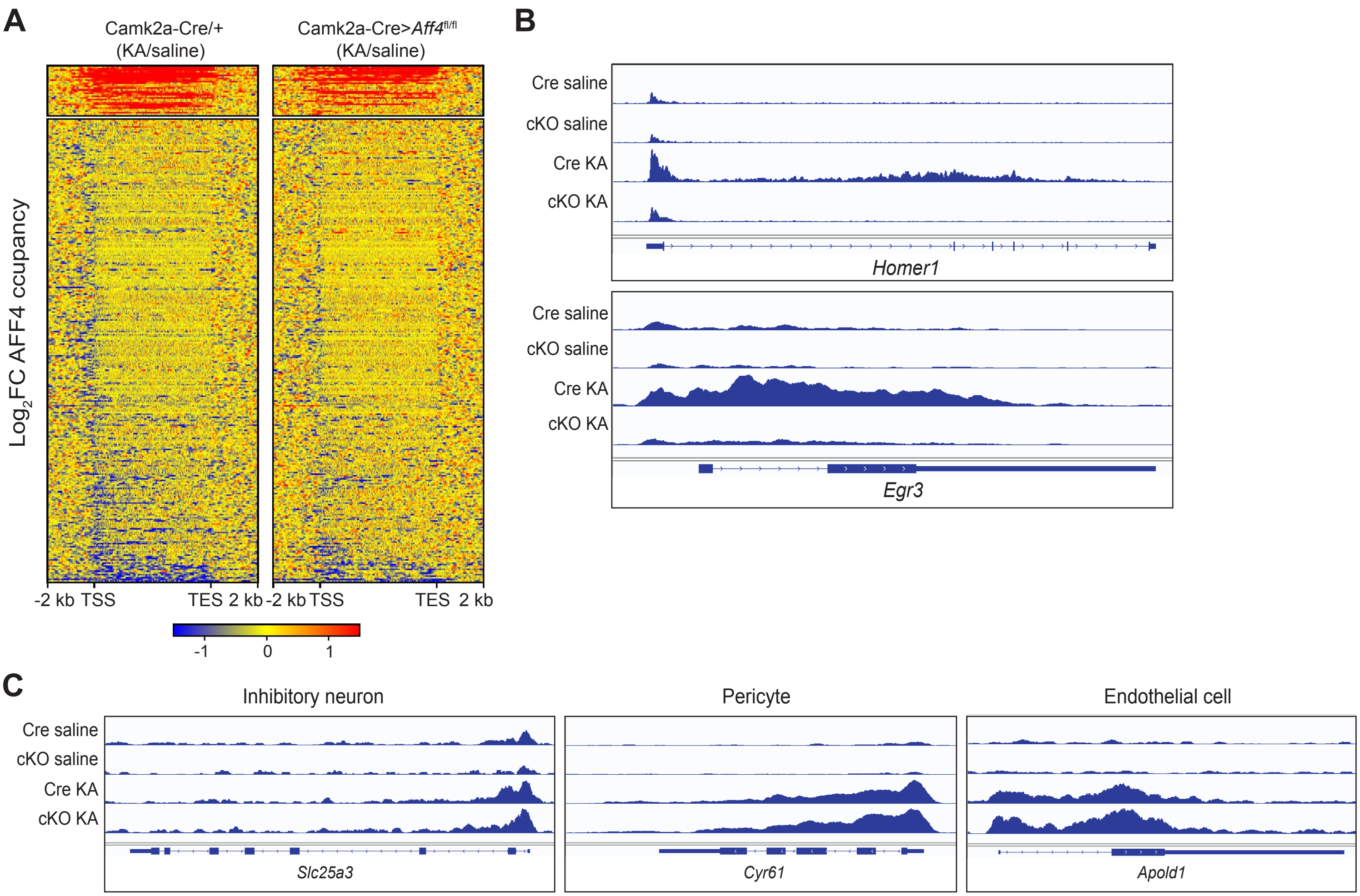

##### Fig. S4 Sonn et al.

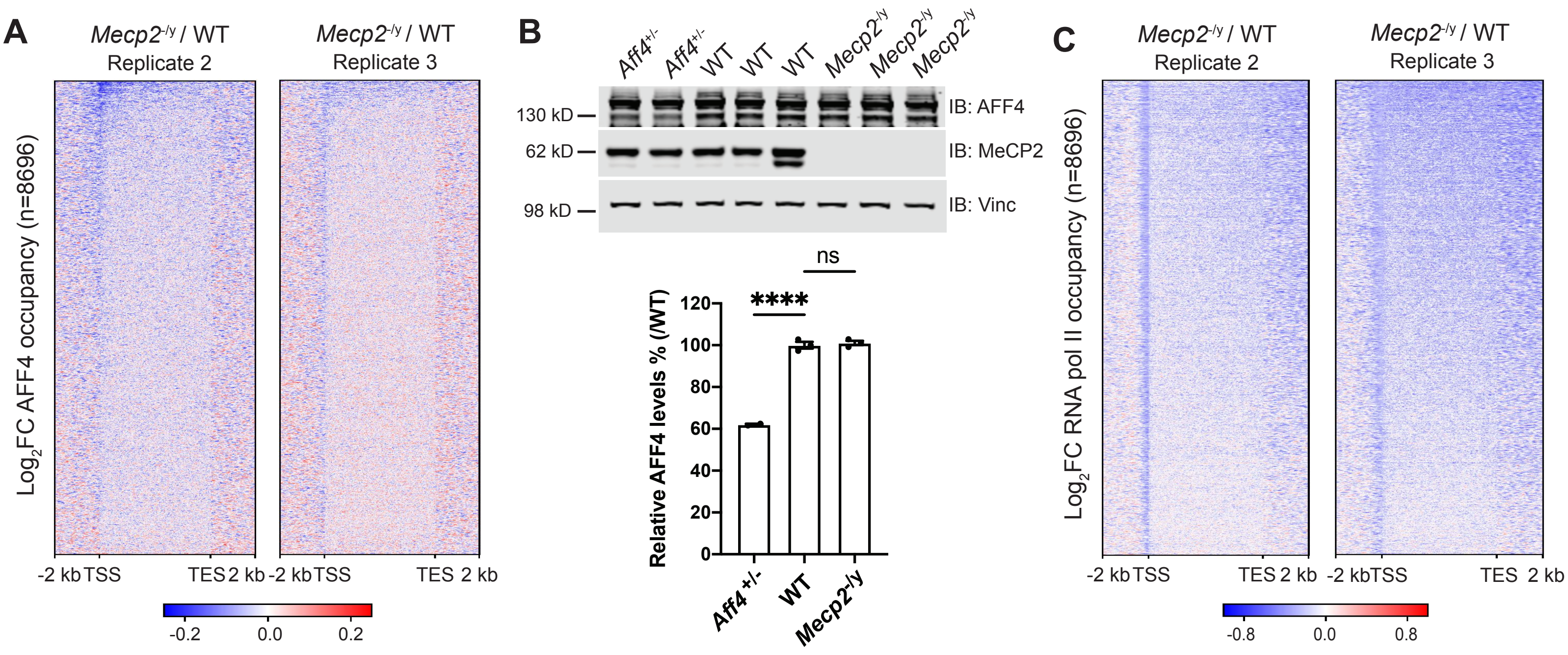

##### Fig. S5 Sonn et al.

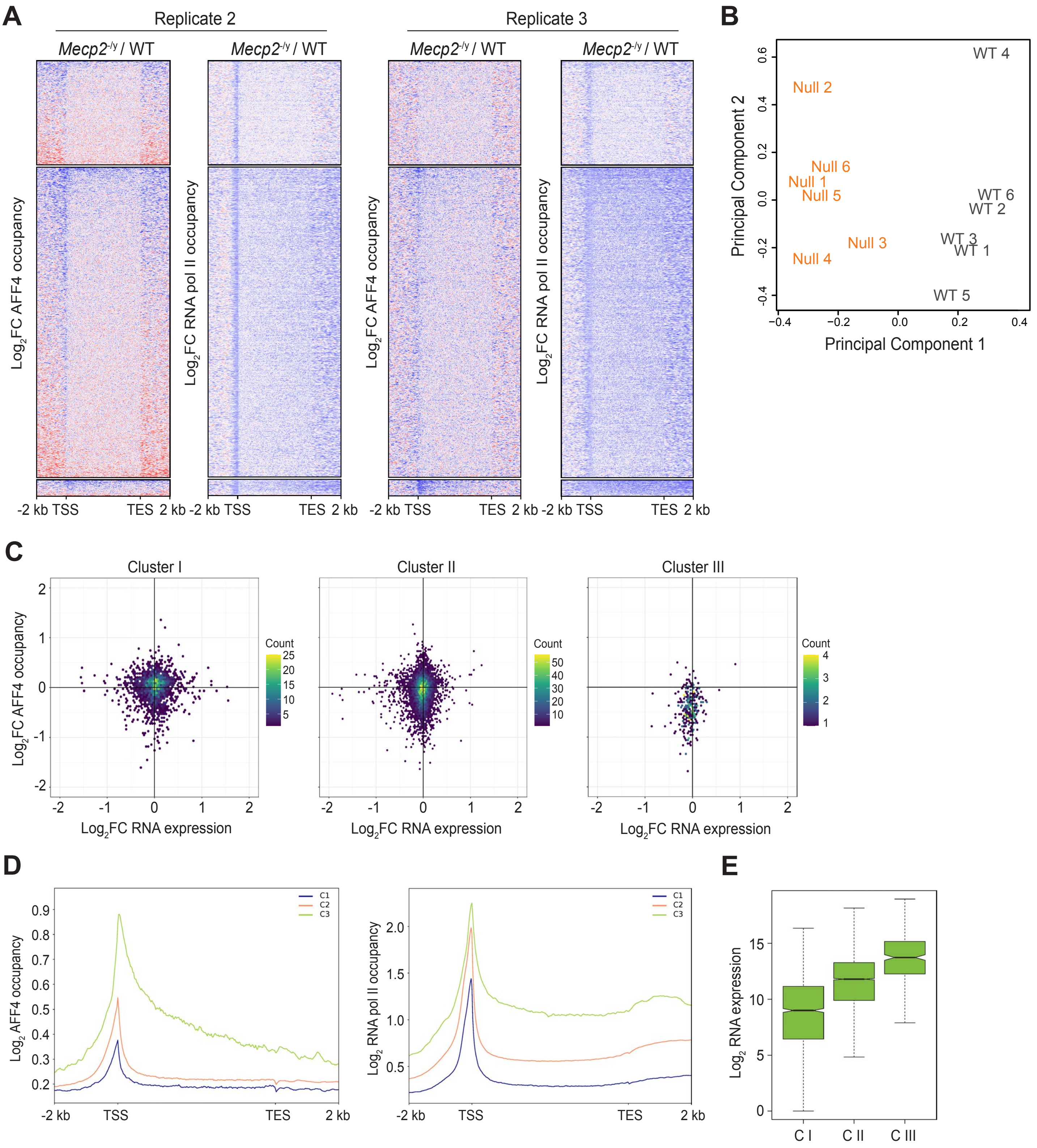
